# Multiple levels of triggered factors and positive effect of cell-to-cell bottleneck in mutation repair of RNA viruses

**DOI:** 10.1101/2021.06.28.450159

**Authors:** Shanshan Liu, Chengming Yu, Guowei Geng, Chenyu Su, Xuefeng Yuan

## Abstract

Debilitating mutation in RNA viruses can be repaired via different mechanisms, while triggering factors of mutation repair were poorly understood. In this study, multiple levels of triggering factors of mutation repair was identified based on genetic damage of tRNA-like structure (TLS) in cucumber mosaic virus (CMV). TLS mutation in different RNAs of CMV distinctively impacted the pathogenicity and mutation repair. Relative quantity defect of RNA2 or quality defect of RNA3 resulting from TLS mutation was correlated with high rate of mutation repair, and TLS mutation of RNA1 failed to be repaired. However, TLS mutation of RNA1 can be repaired in the mixed inoculation with RNA2 having pre-termination mutation of 2b or at the low dose of original inoculation, especially around dilution end-point. Taken together, TLS mutation resulting into quality or quantity defect of viral genome or TLS mutation at low dose around dilution end-point was inclined to be repaired. In addition, different levels of mutation repair of TLS necessarily required the cell-to-cell movement, which implied the positive effect of cell-to-cell bottleneck on evolution of low-fitness virus, a phenomenon opposite to the Muller ratchet. This study provided important revelations on virus evolution and application of viral mild vaccine.

**Author summary:** Due to the low-fidelity of replicase, debilitating RNA viruses can be repaired through different mechanisms, which implied the resilience of RNA viruses. In this study, we identified multiple levels of triggered factors and occurrence occasion of mutation repair using the divided genome of CMV, which contained conserved cis-element tRNA-like structure (TLS) at the 3’end. TLS mutation of different RNA in CMV presented different rate of mutation repair from 0-80%. TLS mutation resulting into genomic quality or quantity defect or at low dose around dilution end-point was inclined to be repaired. However, all above types of mutation repair necessarily required cell-to-cell movement, which presented the positive effect of cell-to-cell bottleneck on virus evolution and increased fitness of low-fitness RNA viruses. It is an opposite phenomenon to the Muller ratchet, in which bottleneck always decreased the fitness of viruses. Except to identify the triggering factors of mutation repair in RNA viruses, this study also provided important revelations on creation and application of mild vaccine based on RNA viruses.

## Introduction

Many factors caused different type of mutations in RNA viruses, which have evolved several tools to maintain their genome integrity due to the infidelity of the replicative machinery (Agol and Gmyl, 2018). Negative selection can eliminate less-fit or lethal variants, while some debilitating mutations in RNA virus can be repaired or remodeled by reversion or compensatory mutation through RNA recombination even in trans complementation from other coinfecting viruses (Barr and Fearns, 2010; Agol and Gmyl, 2018). RNA Recombination played essential role on the genetic evolution as well as genetic stability of all living organism, which is responsible for the rearrangement of viral genes, the repair of debilitating mutations, and the acquisition of nonself sequences (Sztuba-Solińska *et al*., 2011). Based on the infidelity of the replicative machinery, genome of well-adapted RNA viruses can maintain their identity, whereas genome of weak-adapted or debilitated RNA viruses are unstable and can be repaired even become higher-fitness variant (Agol and Gmyl, 2018). RNA recombination is one of the strongest forces shaping robustness and resilience of RNA viruses, which are two connected aspects of viral evolution, although robustness and resilience of RNA viruses seems to be conflicative (Sztuba-Solińska *et al*., 2011; Agol and Gmyl, 2018).

Two models have been proposed to interpret RNA recombination: the potential cleavage-religation of precursor RNAs during the nonhomologous recombination and the replicase-mediated copy-choice mechanism during both homologous and nonhomologous recombination (Chetverin *et al*., 1997; Nagy *et al*., 1998; Galleiet *et al*., 2004; Gmyl and Agol, 2005). The latter is the most widely accepted model for recombination. Replicative-related RNA recombination is mediated by copy-choice mechanism, which occurs at specific crossover sites between different RNA templates possibly involving in related viruses, distantly related viruses and even host RNAs (Figlerowicz and Nagy, 1997; Sztuba-Solińska *et al*., 2011). In copy-choice mechanism of RNA recombination, the specific crossover site between two RNA templates and error-prone replicase are substrate basis, which ensures the potential occurrence of RNA recombination (Sztuba-Solińska *et al*., 2011). Each RNA virus seems to be capable of recombining since RNA recombination does not require complicated machinery. RNA recombination on different RNA viruses will promote viral evolution, while RNA recombination on debilitating viral genome may cause mutation repair. However, factors triggering mutation repair or determining the frequency of mutation repair via RNA recombination was unclear.

In order to evaluate the factor triggering occurrence and frequency of mutation repair mediated by RNA recombination, deleterious mutation and the specific crossover site among different RNAs are required simultaneously. In this study, the divided genome of cucumber mosaic virus (CMV) was used to evaluate the relationship between debilitating genome and mutation repair mediated by RNA recombination, because RNA1, RNA2 and RNA3 of CMV have identical 3’UTR containing conserved tRNA-like structure (TLS), which was essential for replication and had some core cis-elements such as caacg loop (Sivakumaran *et al*., 2000). Identical mutations on TLS in RNA1, RNA2 or RNA3 presented quite different frequency of mutation repair via RNA recombination. Further assay suggested that mutation repair can be triggered by not only quality or quantity defect of genome caused by mutation but also debilitating genome at low original dose. In addition, all these types of mutation repair on TLS mutation required cell-to-cell movement. These results provided new revelation on viral evolution and mild vaccine application.

## Results

### TLS mutation in different RNAs of CMV presented remarkable difference of repair frequency mediated by copy-choice type of RNA recombination

In order to identify the relationship between mutation repair and debilitating genome, debilitating genome was required. In this study, the tRNA-like structure (TLS) was mutated to cause debilitating genome because TLS was previously reported to be essential for in vitro replication of CMV (Sivakumaran *et al*., 2000). TLS of RNA1, RNA2 or RNA3 was respectively mutated in two ways, which included replacement of caacg loop by 18 nt sequences and insertion of 18 nt sequences in caacg loop (Fig. 1A & Supplementary table 1). At 14dpi of agroinfiltration at OD_600_=1.2, viral progeny RNAs in different plants were detected by RT-PCR and DNA sequencing to identify the characteristic of mutation sites. Two type of mutation on TLS in RNA1 (R1-T_m1_ and R1-T_m2_) still existed in progeny RNAs and did not appear mutation repair, while mutation of TLS in RNA2 or RNA3 presented different frequency of mutation repair in progeny RNAs. Mutation repair frequency of R2-T_m1_, R2-T_m2_, R3-T_m1_ and R3-T_m2_ was 50%, 37.5%, 80% and 62.5%, respectively (Fig. 1B). Same type of TLS mutation in RNA3 had higher mutation repair frequency than that in RNA2. In addition, replacement type mutation on caacg loop (m1) had higher mutation repair than insertion type mutation on caacg loop (m2). Taken together, identical TLS mutation in RNA1, RNA2 or RNA3 of CMV presented different characteristic of mutation repair, including the sureness of occurrence and the frequency of occurrence.

**Figure 1.**
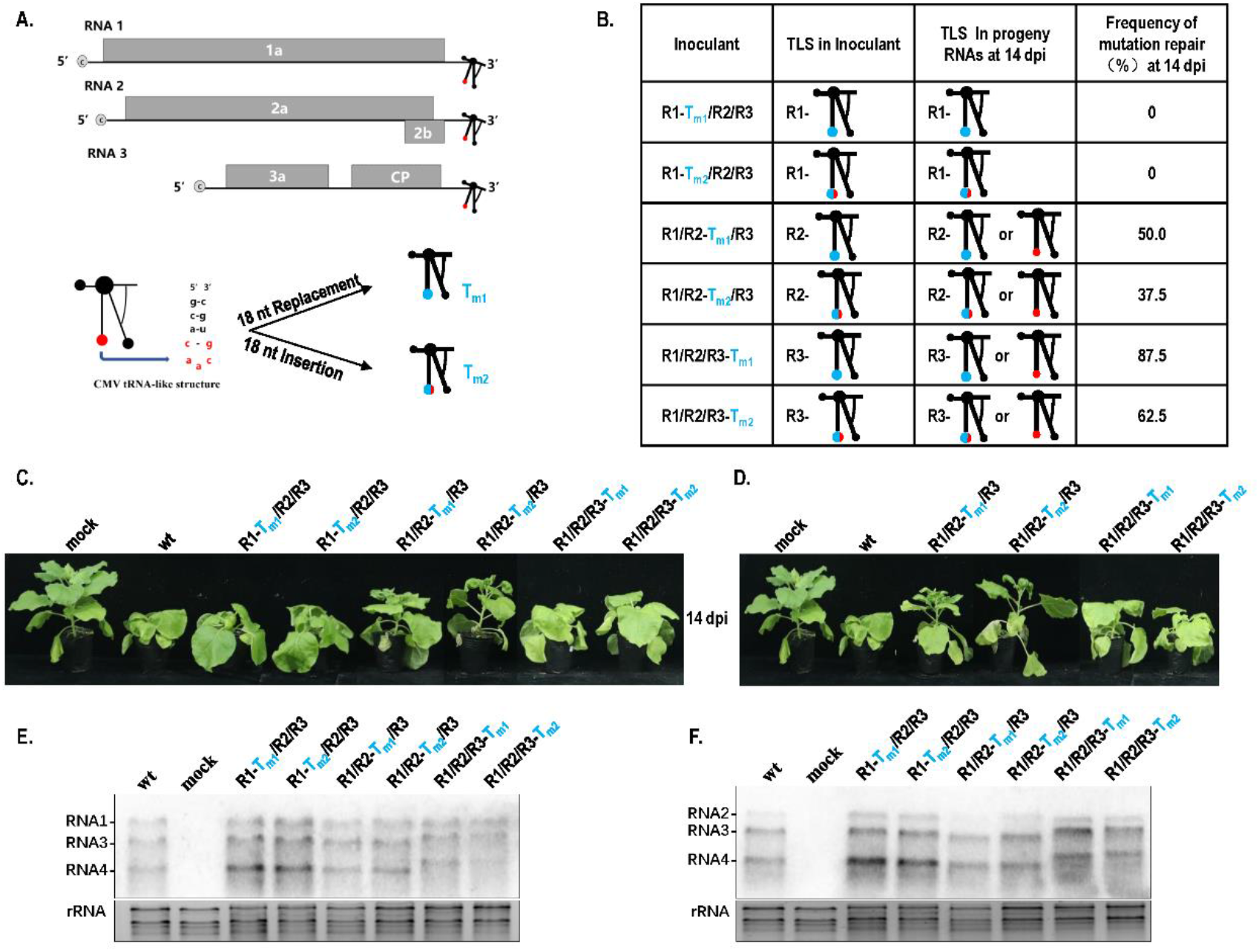
Different mutation repair frequency and pathogenicity of TLS mutation in different RNAs of cucumber mosaic virus. **A. Schematic of cucumber mosaic virus (CMV) and mutation on tRNA-like structure (TLS)**. TLS was indicated by special branch structure and essential caacg loop was indicated by solid red circle. Mutation of 18 nt replacement on caacg (Tm1) or 18 nt insertion before caacg (Tm2) was also indicated by solid blue cricle. **B. Frequency of mutation repair of TLS mutation in RNA1, RNA2 or RNA3 of CMVFny**. R1, RNA1 of CMVFny; R2, RNA2 of CMVFny; R3, RNA3 of CMVFny. **C. Pathogenicity of different TLS mutants without mutation repair**. wt, RNA1/RNA2/RNA3 of wild type CMVFny. **D. Pathogenicity of different TLS mutation with mutation repair**. **E. Northern blot for different TLS mutation without mutation repair using probes against RNA1, RNA3 and RNA4**. **F. Northern blot for different TLS mutation without mutation repair using probes against RNA2, RNA3 and RNA4**.

Due to the conservation of TLS region among RNA1, RNA2 and RNA3 in CMV, RNA recombination may be the candidate mechanism to mediate mutation repair on TLS mutation. To identify whether RNA recombination accomplished the mutation repair on TLS mutation in CMV, identical TLS mutant of RNA1, RNA2 and RNA3 were mixed to simultaneously infect plants followed by analysis of mutation repair (Supplementary Fig. 1). Mixed infection of R1-T_m1_/R2-T_m1_/R3-T_m1_ or R1-T_m2_/R2-T_m2_/R3-T_m2_ presented slight pathogenicity without stunting (Supplementary Fig. 1A). These two types of mixed infection did not present mutation repair of TLS mutation (Supplementary Fig. 1B). It is suggested that mutation repair of TLS mutation was mediated by copy-choice type of RNA recombination. R1-T_m1_/R2-T_m1_/R3-T_m1_ or R1-T_m2_/R2-T_m2_/R3-T_m2_ failed to be repaired, which was due to the absence of copy-choice template through RNA recombination.

### Quality or quantity defect of genomic RNAs resulting from TLS mutation in RNA2 or RNA3 of CMV triggered mutation repair

Based on above data, single TLS mutation of RNA2 or RNA3 can be repaired, while identical TLS mutation of RNA1 failed to be repaired (Fig. 1A & 1B). For TLS mutation of RNA1, RNA2 or RNA3, the prerequisite of RNA recombination was possessed, but occurrence frequency was remarkable difference. It is implied that mutation repair of TLS mutation mediated by RNA recombination was triggered by not only own characteristic of TLS mutation, but also other factors. To identify why TLS mutation of different RNAs in CMV had different rate of mutation repair, the effect of different TLS mutation on CMV pathogenicity and RNA accumulation was analyzed. To exclude the obstruction of repaired mutation, we selected the special samples, in which TLS mutation was not repaired (Fig. 1C). Mutant of R1-T_m1_ or R1-T_m2_ had similar pathogenicity with wt, which caused severe stunting (Fig. 1C). Mutant of R2-T_m1_ or R2-T_m2_ had weak pathogenicity causing slight stunting (Fig. 1C). Mutant of R3-T_m1_ or R3-T_m2_ had weaker pathogenicity than wt and presented weaker stunting than wt (Fig. 1C). Identical TLS mutation in RNA1, RNA2 or RNA3 had different effect on CMV pathogenicity. When TLS mutation in RNA2 or RNA3 was repaired to wt, symptom still had similar pattern with mutant precursor (Fig. 1D), which implied that mutation repair required a process, not a sudden action. To identify why TLS mutation in RNA1, RNA2 or RNA3 had different effect on CMV pathogenicity, characteristic of different RNAs of CMV were analyzed by Northern blotting. Based on the signal intensity of Northern blot, single TLS mutation in RNA1, RNA2 or RNA3 did not affect the RNA accumulation level of total CMV genome (Fig. 1E&1F). However, single TLS mutation in RNA1, RNA2 or RNA3 had different effect on special RNA segment when relative ratio between RNAs was analyzed. TLS mutation of R1-T_m1_ or R1-T_m2_ did not affect relative synthesis of RNA1 as well as RNA2 and RNA3 (Fig. 1E &1F). Mutant of R2-T_m1_ almost dismissed the synthesis of RNA2, while R2-T_m2_ still had RNA2 synthesis with some relative reduction (Fig. 1F). Mutant of R3-T_m1_ or R3-T_m2_ unusually increased the size of RNA3 as well as its subgenomic RNA (RNA4) (Fig. 1E & 1F). Although TLS was conserved in different RNAs of CMV, TLS mutation in RNA1, RNA2 or RNA3 had entirely different effect on synthesis of corresponding RNA (Fig. 1E & 1F). Taken together, repair frequency of TLS mutations in different RNAs of CMV were correlated with their effect on CMV pathogenicity and synthesis of corresponding RNAs (Fig. 1). Quality defect of RNA3 size caused by TLS mutation (T_m1_ or T_m2_) in RNA3 triggered high frequency of mutation repair at 80% or 62.5%. Quantity defect of RNA2 synthesis caused by TLS mutation (T_m1_ or T_m2_) in RNA2 also triggered high frequency of mutation repair at 50% or 37.5%. TLS mutation in RNA1 did not cause quantity or quality defect, which is correlated with the nonoccurrence of mutation repair. It is suggested that quality or quantity defect of genome segment of CMV caused by TLS mutation was the important triggered factor for mutation repair.

### Mutation repair of debilitating genome can be triggered by critical threshold low dose, especially around dilution end-point

Although TLS mutation of RNA1 failed to be repaired (Fig. 1), after all it caused the debilitating TLS in RNA1. Whether can the debilitating TLS in RNA1 be repaired at other special condition such as mixed infection with other type of mutant causing genome defect? Pre-termination mutant (R2-2bPT) of gene-silencing suppressor 2b presented mild pathogenicity without stunting and low RNA accumulation (Fig. 2). When TLS mutation in RNA1 (R1-T_m1_ or R1-T_m2_) and R2-2bPT co-infected plants, TLS mutant in RNA1 was repaired at the rate of 100% in progeny RNAs (Fig. 2B), while single TLS mutation in RNA1 failed to be repaired (Fig. 1B). Similarly, when TLS mutation in RNA3 (R3-T_m1_ or R3-T_m2_) and R2-2bPT co-infected plants, TLS mutation was repaired at the rate of 100% in progeny RNAs (Fig. 2B), which was higher than that of single TLS mutation (R3-T_m1_ or R3-T_m2_) (Fig. 1B). Presence of R2-2bPT can remarkably improve the repair frequency of TLS mutation in RNA 1 or RNA3, which may be due to relative low dose of RNA accumulation resulting from the defect of gene-silencing suppressor 2b (Fig. 2C). To further confirm the relationship of the low dose of CMV genome and mutation repair of TLS mutation, dilution inoculation of R1-T_m1_ /R2/R3 or R1-T_m2_ /R2/R3 was performed (Fig. 3). Mutation repair of TLS mutation in RNA1 appeared at the condition of OD_600_=1.2*10^−4^ (Fig. 3A & 3B), which was exactly the dilution end-point of wile type CMV (Fig. 3C). In addition, wild type of CMV or CMV with debilitating cis-element had the similar level of dilution end-point, because there was no CMV accumulation at the inoculation of OD_600_=1.2*10^−5^ and OD_600_=1.2*10^−6^ (Fig. 3). It is suggested that critical threshold point of low dose of viral genome, especially around dilution end-point can trigger mutation repair of debilitating TLS in RNA1, although the debilitating TLS in RNA1 did not present negative effect on pathogenicity and RNA accumulation of CMV (Fig. 1 & Fig. 3).

**Figure 2.**
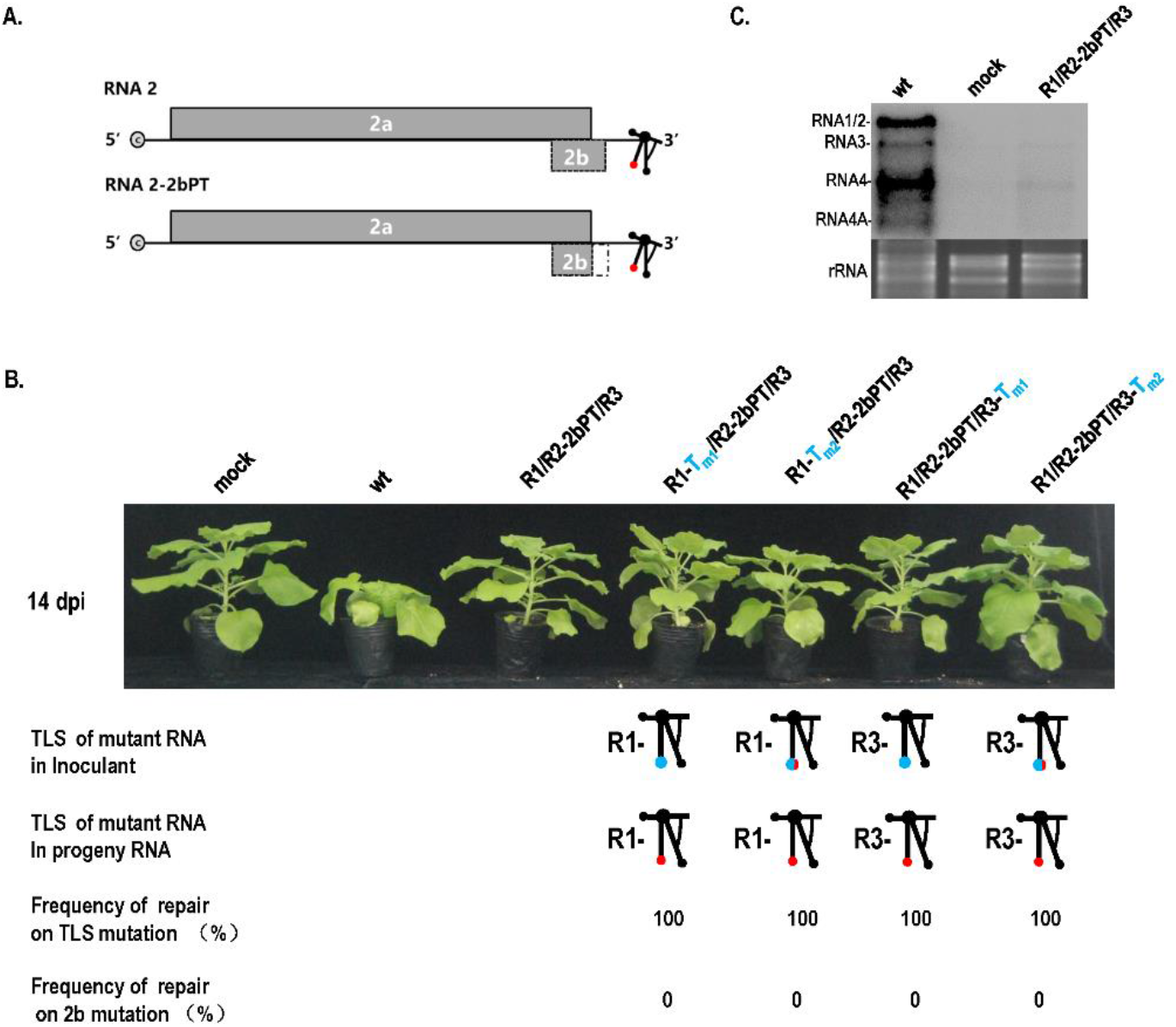
Mutation repair on TLS mutation of RNA1 or RNA3 in the presence of 2b pre-termination in RNA2. **A. Schematic of 2b pre-termination mutant in RNA2**. Note: dotted rectangle indicated the deletion of partial 2b ORF caused by pre-termination of 2b. PT, pre-termination. **B. Mutation repair on TLS mutation of RNA1 or RNA3 in the presence of 2b-defected RNA2**. wt, RNA1/RNA2/RNA3 of wild type CMV_Fny_. R2-2bPT, mutant of 2b pre-termination. **C. Effect of 2b pre-termination on RNA accumulation of CMV**.

**Figure 3.**
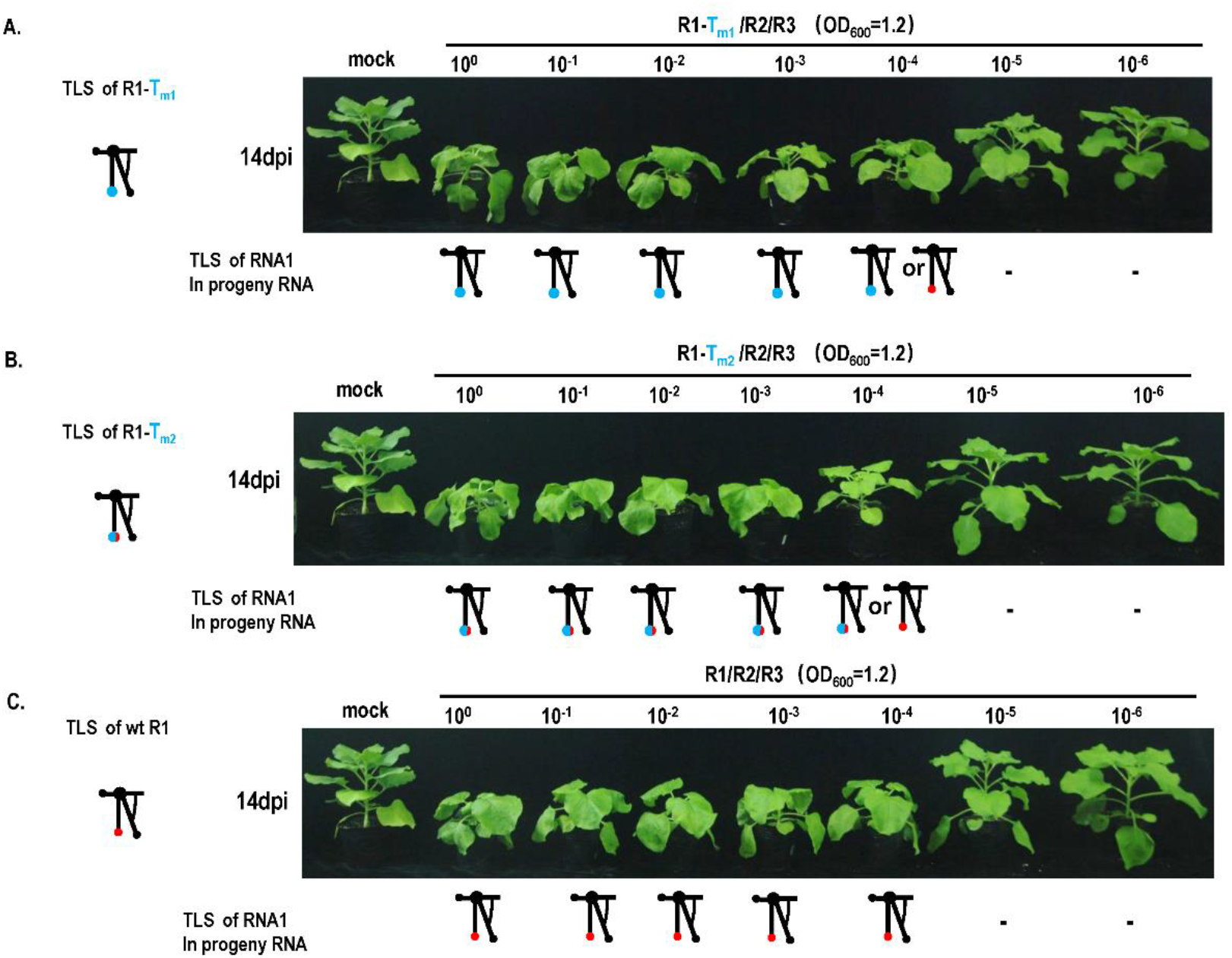
Dilution inoculation of CMV_Fny_ or its variants and mutant repair assay. **A. Mutation repair assay under dilution inoculation of R1-Tm1/R2/R3**. R1, RNA1 of CMV_Fny_; R2, RNA2 of CMV_Fny_; R3, RNA3 of CMV_Fny_;-,no detectable viral RNA. **B. Mutation repair assay under dilution inoculation of R1-Tm2/R2/R3**. **C. Mutation repair assay under dilution inoculation of CMV**_**Fny**_.

### Cell-to-cell movement was necessarily required for mutation repair of TLS mutation, which presented the positive effect of bottleneck on mutation repair

Based on above data, mutation repair can be triggered by quality or quantity defect of genome resulted from TLS mutation or TLS mutation at the special low dose of original inoculation, which was related with low-fitness of viral genome (Fig. 1&Fig. 2&Fig. 3). In addition to these multiple levels of trigger factors, occurrence of mutation repair may require special occasion. Previous study had reported that some mutation of essential element was not repaired in protoplast system but repaired directly or indirectly in plants (Yuan *et al*., 2010). It is implied that the occurrence of mutation repair may require the cell-to-cell movement of virus. In order to analyze the role of cell-to-cell movement on mutation repair, cell-to-cell movement-defected mutant R3-m5 was constructed via two Ala replacement corresponding to position 549-554 in 3a ORF (Ding *et al*., 1995; Fig. 4A&4B). When R2-T_m1_ or R2-T_m2_ and R3-m5 were co-inoculated into plants, mutation repair failed to be occurred (Fig. 4C), while single R2-T_m1_ or R2-T_m2_ had high rate of mutation repair at 50% or 37.5% (Fig. 1B). When R1-T_m1_ or R1-T_m2_ and R3-m5 were co-inoculated into plants, mutation repair failed to be occurred at any dose (Fig. 5), while single R1-T_m1_ or R1-T_m2_ appeared mutation repair around dilution end-point (Fig. 3). It is suggested that mutation repair of TLS mutation required the cell-to-cell movement of virus.

**Figure 4.**
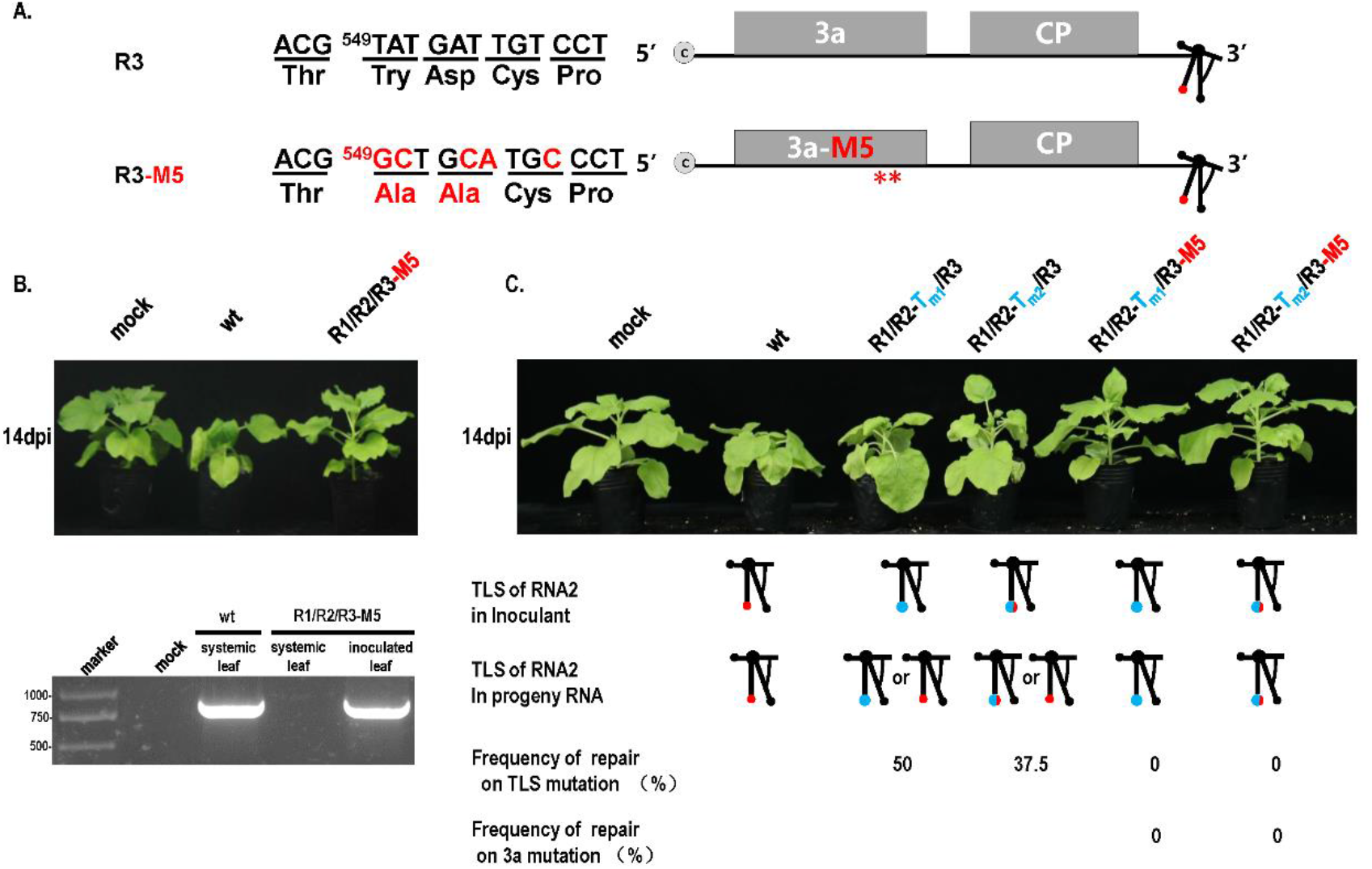
Repair assay of TLS mutation in RNA2 with the movement-defected CMV. **A. Mutation on movement protein 3a in CMV**. Mutation of nucleotide and amino acid was red color. **B. Effect of 3a mutation on pathogenicity of CMV**. **C. Effect of 3a mutation on mutation repair of TLS mutation in RNA2**.

**Figure 5.**
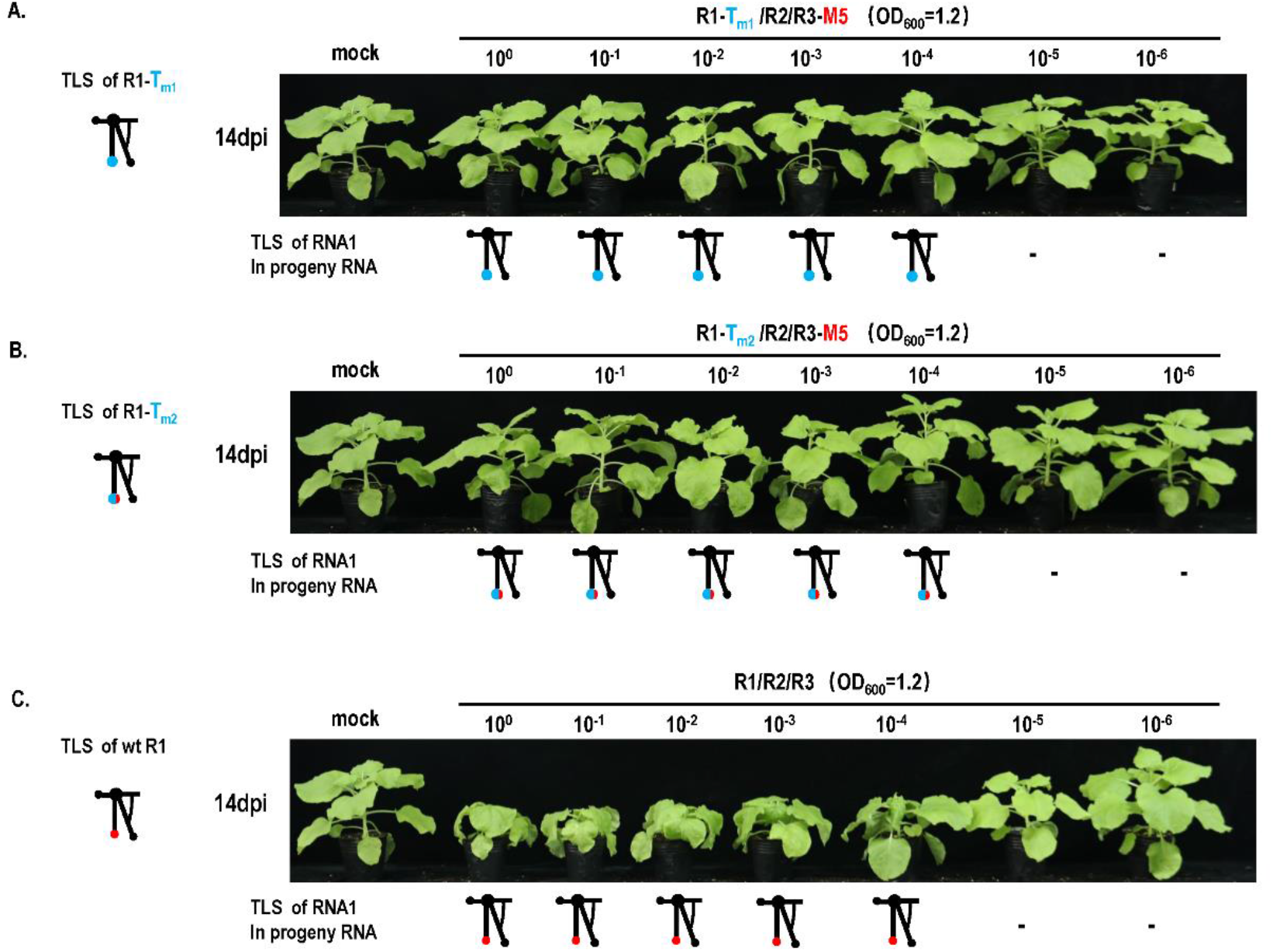
Effect of cell-to-cell movement on mutant repair in dilution inoculation assay. **A. Mutation repair assay under dilution inoculation of R1-Tm1/R2/R3-m5**. R1, RNA1 of CMV_Fny_; R2, RNA2 of CMV_Fny_; R3-m5, a3 mutation in RNA3 of CMV_Fny_. **B. Mutation repair assay under dilution inoculation of R1-Tm2/R2/R3-m5**. **C. Mutation repair assay under dilution inoculation of wild type CMV**.

Taken together, mutation repair of TLS mutation in CMV presented multiple levels of triggered factors and obligated requirement of cell-to-cell movement of virus (Fig. 6). Multiple levels of triggered factors in mutation repair included mutation characteristic of TLS, quantity defect or quality defect of genome segments, or quantity defect of whole genome, especially around dilution end-point. Obligated requirement of cell-to-cell for mutation repair presented the positive effect of bottleneck on mutation repair, which was an opposite phenomenon to the Muller rachet (Fig. 6).

**Figure 6.**
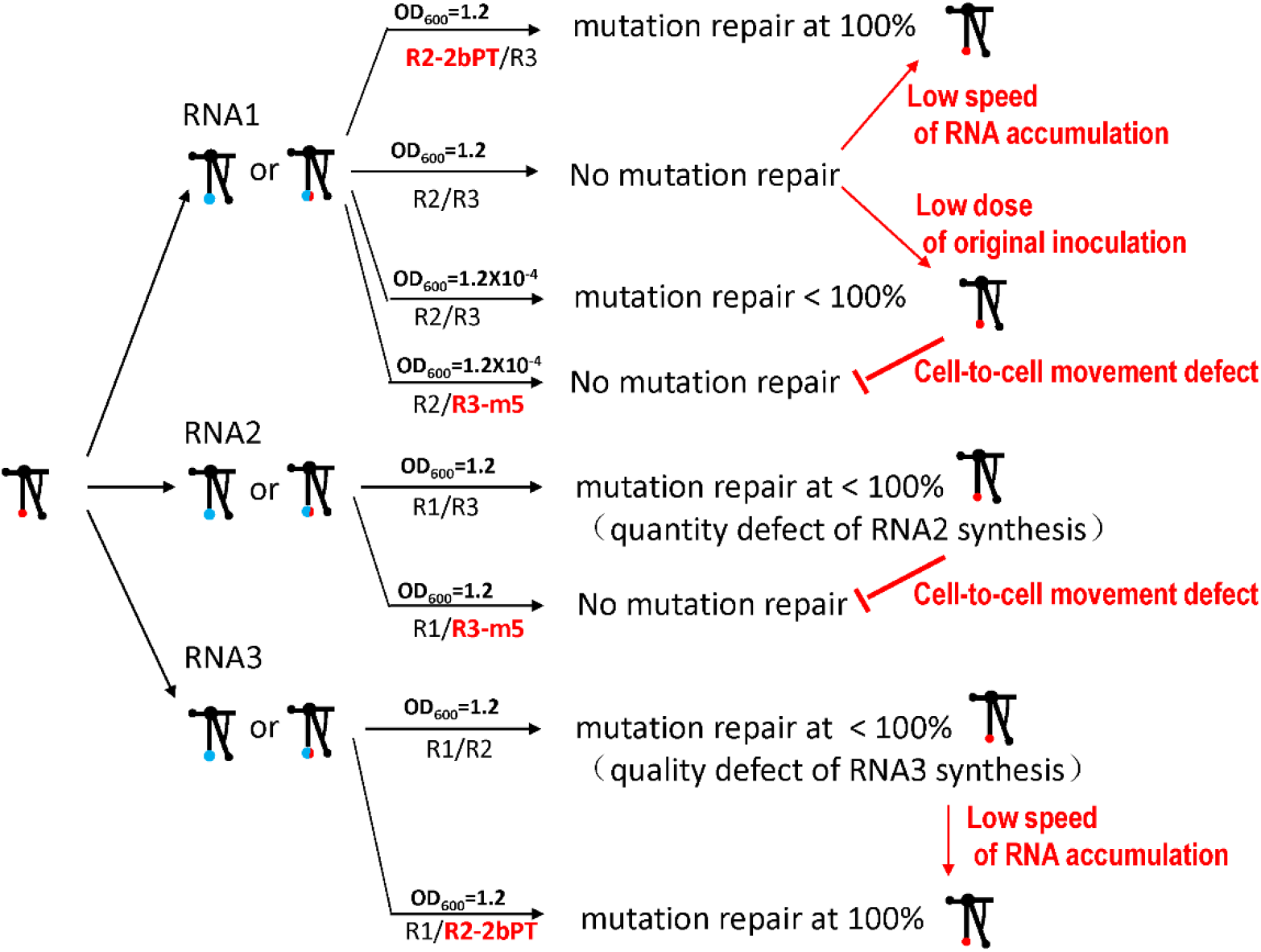
Summary of mutation repair on TLS mutation in different RNAs of CMV. Note: Red arrows indicated the promotion on mutation repair; red vertical lines indicated the inhibition on mutation repair.

## Discussion

### Identical TLS element in different RNAs of CMV played different role on pathogenicity in CMV

TLS was the essential element for *in vitro* replication of CMV (Sivakumaran *et al*., 2000). Some reports also confirmed the importance of TLS on *in vivo* RNA accumulation and pathogenicity of CMV as well as other viruses (Boccard and Baulcombe, 1993; Chapman *et al*., 1998; Dreher, 2009; Rao and Kao, 2015). TLS was conserved among RNA1, RNA2 and RNA3 of CMV, however no previous reports compared the role of TLS on *in vivo* synthesis of RNA1, RNA2 or RNA3 of CMV. In this study, identical TLS mutation in RNA1, RNA2 or RNA3 presented remarkable different effect on synthesis of corresponding RNA as well as different frequency of mutation repair. TLS mutation in RNA1 did not affect synthesis of RNA1 and failed to be repaired at high dose of original inoculation. TLS mutation in RNA2 caused the reduction of RNA2 synthesis and had high frequency of mutation repair. TLS mutation in RNA3 caused unusual change of RNA3 size and had highest frequency of mutation repair (Fig. 1). All these data illuminated that TLS of RNA1, RNA2 or RNA3 played different role on corresponding RNA. It is suggested that TLS did not independently play role on synthesis of corresponding RNAs in vivo and different RNAs in CMV may contain other elements interacting with TLS to regulate synthesis of corresponding RNAs. This potential interaction with TLS may or may not compensate the effect of mutation of TLS, so TLS mutation of different RNAs had different frequency of mutation repair. Taken together, Identical TLS element in different RNAs of CMV played different role on pathogenicity in CMV.

### Multiple levels of factors can trigger the mutation repair

Error-prone performance of the viral RdRp was inclined to cause genetic mutation such as point mutation, insertion, deletion and replacement during life cycle (Cooper 1968; Deffit and Hundley, 2016; Neme *et al*., 2017). In addition, other adverse factors or engineered genetic operation can also produce mutation on viral genome (Barr and Fearns, 2010). All these natural or artificial mutation could have several effects on virus genome including neutral effect, debilitating effect and lethal effect. Viruses have evolved numbers of mechanism to deal with these mutations in two directions including negative selection on less-fit variants and rehabilitation on mutation directly or indirectly (Agol and Gmyl, 2018). The accomplishment of the latter was relied on the infidelity of viral itself RdRP through RNA recombination or complementation by other coinfecting viruses (Gmyl and Agol, 2005; Barr and Fearns, 2010; Shirogane *et al*., 2013). Based on the data in this study and some previous reports, mutation repair was related with three levels of trigger factors. For the first level, mutation sites appeared on essential cis-elements such as TLS. Tons of previous report and this study reported mutation repair in this level (Bujarski and Kaesberg, 1986; Rao *et al*.,1989; Hema *et al*., 2005; Rao, 2006). For the second level, mutation caused quantity or quality lack of genome segments such as RNA2 or RNA3 in CMV. For the third level, mutated genome at low original dose around dilution end-points. All these levels of factors caused low-fitness of virus, which was inclined to trigger mutation repair.

### Positive effect of cell-to-cell movement for mutation repair implied the positive effect of bottleneck on virus evolution

In addition to the existence of above triggered factors, occurrence of mutation repair of TLS mutation necessarily required cell-to-cell movement, which implied the positive and exclusive effect of bottleneck on mutation repair. However, previous studies have reported that bottleneck events are generally disadvantageous for viral population (Chao,1990; Duarte *et al*., 1992; Yuste *et al*., 1999; Lázaro *et al*., 2003; de la Iglesia and Elena, 2007; Jaramillo *et al*., 2013), which was mainly due to the reduction of the genetic variation in virus population and fitness by various bottlenecks, a phenomenon known as the Muller ratchet (Manrubia *et al*., 2005; Zwart and Elena, 2015; Poirier and Vignuzzi, 2017; Huang *et al*., 2017; Clarke *et al*., 1993; Escarmís *et al*., 2006; Escarmís *et al*., 2009). The possible positive effect of bottleneck was the removal of defective interfering viruses and selection on evolutionary trajectories in rugged fitness landscapes (Zwart *et al*., 2008; Wright, 1932; Elena *et al*., 2010; Lalić and Elena, 2012; Silva and Wyatt, 2014). In this study, positive effect of bottleneck on mutation repair identified an opposite phenomenon to the Muller ratchet and provided new insight on evolution. So, the effect of bottleneck on virus evolution should be considered from two sides. For well-adapted or wild type of virus, bottleneck mainly presented disadvantage resulting into the reduction of the genetic variation of virus population and fitness after overcoming barriers, the most case in the Muller ratchet. For weak-adapted or debilitating virus, bottleneck presented advantage shown as the obligated requirement of cell-to-cell movement for mutation repair in this study. Without the cell-to-cell movement, debilitating mutation failed to be repaired, which was similar with the case in single-cell system such as protoplast. Through the cell-to-cell barrier, TLS mutation was inclined to be repaired and virus fitness was improved. It is suggested that bottleneck may present different effect on virus with different fitness.

### Revelation on construction and application of mild vaccine from mutation repair

Priority characteristic of mild vaccine was the safety, which require the stability of mutation causing mild pathogenicity (Ziebell and Carr, 2010; Hasiów-Jaroszewska *et al*., 2011; Ziebell and MacDiarmid, 2017). However, mutation repair frequently occurred due to the low fidelity of RdRp in viruses, which was the biggest threaten on the safety of mild vaccine (Agol and Gmyl, 2018). Analysis on trigger factor and occurrence occasion of mutation repair in this study provided important revelation on construction and application of mild vaccine to avoid the mutation repair in any possible way. The original inoculation of mild vaccine should be performed at the high dose, because the original inoculation at the low dose greatly increased the frequency of mutation repair from 0% to 100% such as the case of TLS mutation in CMV RNA1 (Fig. 1& Fig. 3). There may appear the low dose around dilution end-point when mild vaccine entered a new plant during the transmission due to the low acquisition and secondary infection of mild vaccine by aphid. This special low dose may be inclined to trigger mutation repair, which will produce wild type of CMV destroying the safety of mild vaccine. Therefore, transmission characteristic of mild vaccine should be removed to avoid the potential mutation repair.

### Materials and methods Plasmid construction

All mutant plasmids were derived from pCB301-Fny1, pCB301-Fny2 and pCB301-Fny3, which respectively contained wild type of RNA1 (D00356), RNA2 (D00355) and RNA3 (D10538) sequences of CMV_Fny_ isolate (Yao *et al*., 2011). Two types of mutants on the TLS were respectively constructed in RNA1, RNA2 or RNA3 of CMV_Fny_. Mutant T_m1_ of TLS was the replacement of essential caacg loop in TLS by new 18 nt sequences ggatccactagtcccggg. Mutant T_m2_ of TLS was the insertion of new 18 nt sequences ggatccactagtcccggg before caacg loop in TLS. The mutant R2-2bPT was the insertion of two stop codons (UAAUAG) and MCS (ggatccactagtcccggg) after position ^2661^G in RNA2 to cause the pre-termination of 2b protein. Mutant R3-m5 was the mutation of ^549^tatgattgt to ^549^<u>gc</u>tg<u>ca</u>tg<u>c</u> in 3a protein, causing cell-to-cell movement defect.

All above mutants were constructed from the plasmid pCB301-Fny1, pCB301-Fny2 or pCB301-Fny3 through oligo-mediated mutagenesis, which was performed by site mutagenesis mediated by primers. Detailed information of above plasmids and primers was shown in Supplementary table1 and table 2. All mutants were confirmed by DNA sequencing.

### Plasmid transformation and agroinfiltration

All above plasmids were respectively transformed into *Agrobacterium tumefaciens* GV3101 using the CaCl_2_-mediated freeze-thaw method, described previously (Weigel and Glazebrook, 2006). Single *A. tumefaciens* colony was shaking cultured in LB medium with 50 µg/ml kanamycin and 100µg/ml rifampin at 28 °C until OD_600_=1.0-2.0. After centrifugation, the pellets were resolved to OD_600_=1.2 in infiltration solution [10 mM MgCl_2_, 10 mM 2-(N-morpholino) and 0.15 mM acetosyringone]. GV3101 containing plasmids of RNA1, RNA2 or RNA3 as well as the mutants were equally mixed. The mixture was further diluted in dilution assay based on the requirement of different experiments. The mixture was stood at 28 °C for 3 hours and infiltrated into the 4th and 5th leaves of *N. benthamiana* at the six to seven leaf stage. Plants were grown under the condition of 16 hours of photoperiod and 8 hours of darkness at 25°C, as described previously.

### Mutation repair assay

To evaluate the stability and mutation repair of mutation sites in progeny RNAs, total RNA was extracted from inoculated and systematic leaves at 14dpi after agroinfiltration followed by RT-PCR. Specific primers located at upstream and downstream of corresponding mutation sites were used in RT-PCR. RT-PCR products were purified and directly sequenced using specific primers. Mutation repair assay for each mutant was performed at least 10 individual repeats. Detailed information of primers used to amplify mutation regions were shown in supplementary table2.

### Northern blot

Total RNAs were extracted from leaves at 14 dpi after agroinfiltration and Northern blot was performed as previously described (Yuan *et al*., 2006). cDNA probes were labelled using random primers. Four types of cDNA probes were used to detect different genomic and corresponding subgenomic RNAs. cDNA probe 1 (cP1) corresponding to 2951-3150 of RNA1 was designed to exclusively detect RNA1. cDNA probe 2 (cP2) corresponding to 2548-2730 of RNA2 was designed to exclusively detect RNA2. cDNA probe 3 (cP3) corresponding to 1704-1910 of RNA3 was designed to exclusively detect RNA3. cDNA probe 4 (cP4) corresponding to 3’ end of RNAs in CMV was designed to simultaneously detect RNA1, RNA2 and RNA3 as well as corresponding subgenomic RNAs. Detailed information of primers used to amplify PCR products for making these cDNA probes were shown in Supplementary table 2.

## Acknowledgments

The work was funded and supported by National Natural Science Foundation of China (32072382, 31872638). The funders played no role in study design, data collection, interpretation, or the decision to publish this study.

## Author Contributions

S.L. performed and analyzed most of lab experiment data, and C.Y., G.G., and C.S. performed most of statistical analyses and sequencing data analyses. X.Y. wrote the manuscript. All authors discussed the results and commented on the paper.

